# DNA methylation clocks show slower progression of aging in naked mole-rat queens

**DOI:** 10.1101/2021.03.15.435536

**Authors:** Steve Horvath, Amin Haghani, Nicholas Macoretta, Julia Ablaeva, Joseph A. Zoller, Caesar Z. Li, Joshua Zhang, Masaki Takasugi, Yang Zhao, Elena Rydkina, Zhihui Zhang, Stephan Emmrich, Ken Raj, Andrei Seluanov, Chris G. Faulkes, Vera Gorbunova

## Abstract

Naked mole-rats (NMRs) live an exceptionally long life, appear not to exhibit age-related decline in physiological capacity, and are seemingly resistant to age-related diseases. However, it has been unknown whether NMRs also evade aging according to a primary hallmark of aging: epigenetic changes. To address this question, we generated DNA methylation profiles from 329 tissues from animals of known age, at loci that are highly conserved between mammalian species, using a custom Infinium array (HorvathMammalMethylChip40). We observed strong aging effects on NMR DNA methylation, from which we developed seven highly accurate age estimators (epigenetic clocks) for several tissues (pan-tissue, blood, kidney clock, liver clock, skin clock) and two dual species (human-NMR) clocks. By identifying age-related cytosine methylation that are shared between NMR and humans, but not with the mouse, we identified genes and cellular pathways that impinge on developmental and metabolic processes that are potentially involved in NMR and human longevity. The NMR epigenetic clocks revealed that breeding NMR queens age more slowly than non-breeders, a feature that is also observed in some eusocial insects. CpGs associated with queen status were located near developmental genes and those that are regulated by the *LHX3* transcription factor that controls pituitary development. In summary, our study demonstrates that despite a phenotype of reduced senescence, the NMR ages epigenetically through developmental and metabolic processes, and that NMR queens age more slowly than non-breeders.

## INTRODUCTION

The naked molerat (NMR), *Heterocephalus glaber,* has emerged as a popular model for aging studies due to its exceptional lifespan of over 30 years, despite being of comparable size to a laboratory mouse, with a lifespan of approximately 3 years. The relative ease of breeding NMRs in captivity makes them particularly attractive model for laboratory studies. NMRs live in large subterranean colonies in East Africa. The colonies have an eusocial structure^1^, similar to those of ants and bees. NMR social groups consist of a single breeding queen who mates with one to three males (favorites), while the rest of the colony are non-reproductive and most will never breed. This Hystricomorph rodent has a recorded maximum lifespan of 37 years (or more) and appears to be resistant to many age-related diseases^2–8^ including two leading causes of death in humans: cardiovascular disease and cancer^9,10^. Strikingly, the mortality rate of this rodent defies the Gompertz-Makeham law by not increasing with age^11^. The NMR is proposed to be a “non-aging” mammal as it displays negligible senescence, including minimal age-related decline in physiological capacity as measured by several aging biomarkers ^3,11^.

Epigenetic age, as measured by epigenetic clock is arguably the most accurate estimator of age in numerous mammalian species including humans ^12–14^ Hence, it remains to be ascertained whether the NMR epigenome undergoes epigenetic age-related changes.

Cytosine methylation is one of the best characterized epigenetic modifications that modulates gene activity and chromatin structure. In mammals, DNA methylation plays an important role in multiple biological processes including silencing of transposable elements, regulation of gene expression, genomic imprinting, X-chromosome inactivation, carcinogenesis, and aging^15^ Indeed, it was long observed that the degree of cellular DNA methylation changes with age in humans and many mammalian species^16–18^ Following the development of array-based quantification of methylation of specific CpG positions on the human genome, came the insight that methylation levels of multiple CpGs can be consolidated to develop highly-accurate multivariate age-estimators (epigenetic clocks) for all human tissues^12,19^ For example, the human pan-tissue clock combines the weighted average of DNA methylation levels of 353 CpGs into an age estimate that is referred to as DNA methylation age (DNA methylation age) or epigenetic age^20^ Importantly, the difference between DNA methylation age and chronological age (referred to as “epigenetic age acceleration”) is predictive of all-cause mortality in humans, even after adjusting for a variety of known risk factors^21–25^ The notion that epigenetic age acceleration may be indicative of health status was confirmed by multiple reports that demonstrated its association with a multitude of pathologies and health conditions^21,26–29^ Today, epigenetic clocks are regarded as validated biomarkers of human age and are employed in human clinical trials to measure the effects and efficacy of anti-ageing interventions^12,14,30^ The human epigenetic clocks, however, cannot be applied to non-primate species because of evolutionary genome sequence divergence^20^ Hence, it is necessary to develop *de novo* epigenetic clocks that are specific to species of interest, as has been accomplished for mice^13,31–35^ Here, we present epigenetic clocks for the NMR and demonstrate that despite a phenotype of reduced senescence, NMR, like other mammals, age at the epigenetic level. We also reveal the characteristics of individual methylated loci that correlate strongly with age in different NMR tissues. Beyond ageing, the NMR is also of great interest as a model system for understanding the evolution and maintenance of cooperative breeding and eusociality, and extreme socially-induced suppression of reproduction. Eusociality, which is more common in insects such as bees and ants, is characterised by queens that live much longer than workers. We present evidence that NMR queens exhibit slower epigenetic aging rates and reveal characteristics of CpGs and their proximal genes that relate to queen status.

## RESULTS

### Random forest predictors of tissue and sex

We obtained methylation profiles of 382 DNA samples derived from eleven NMR tissues **(Table 1)**: adipose (n=3), blood (n=92), cerebellum (n=5), cerebral cortex (n=15), heart (n=21), kidney (n=33), liver (n=68), lung (n=19), muscle (n=19), skin (n=105), spleen (n=2). These tissues were obtained from naked mole rats that ranging in age from 0 to 26 years-old. All methylation profiles were generated using a custom Illumina methylation array (HorvathMammalMethylChip40), in which 27,917 out of its 37,492 CpGs mapped to the NMR genome (HetGla_female_1.0.100 genome assembly). Random forest predictors for tissue type and sex based on these methylation profiles led to near-perfect accuracy in predicting tissue type (out-of-bag error rate of zero) and sex (only 1 misclassification out of 382 samples) (Supplementary Figure 1). The remarkable accuracy of these predictors based on methylation profiles makes them valuable to validate platemaps and detect potential human errors that may occur when deriving and analyzing DNA methylation data.

**Table 1.** Tissue samples used in the study.

| **Tissue** | **N** | **No.**<br>**Female** | **Mean**<br>**Age** | **Min.**<br>**Age** | **Max.**<br>**Age** |
| --- | --- | --- | --- | --- | --- |
| **Adipose** | 3 | 2 | 2.19 | 0.67 | 3.6 |
| **Blood** | 92 | 73 | 5.4 | 0.67 | 26 |
| **Cerebellum** | 5 | 3 | 1.54 | 0.67 | 3.35 |
| **Cortex** | 15 | 7 | 2.66 | 0.115 | 10.6 |
| **Heart** | 21 | 8 | 2.01 | 0.0055 | 10.6 |
| **Kidney** | 33 | 11 | 4.7 | 0.0055 | 23 |
| **Liver** | 68 | 22 | 7.21 | 0.115 | 26 |
| **Lung** | 19 | 7 | 2.34 | 0.115 | 10.6 |
| **Muscle** | 19 | 6 | 2.72 | 0.115 | 10.6 |
| **Skin** | 105 | 46 | 3.74 | 0 | 25 |
| **Spleen** | 2 | 1 | 0.115 | 0.115 | 0.115 |

N=Total number of samples per species. Number of females. Age: mean, minimum and maximum.

### Epigenetic clocks

To build methylation-based age estimators, we used penalized regression models (elastic net regression) to regress chronological age (dependent variable) on the 27,917 CpGs (covariates) that map to the NMR genome. To arrive at unbiased estimates of the epigenetic clocks, we performed cross-validation analyses and obtained estimates of age correlation R (Pearson correlation between estimated DNAm age and chronological age), as well as the median absolute error. We generated four epigenetic clocks for NMRs that are optimised for blood, kidney, liver and skin respectively. In addition, we also developed a fifth epigenetic clock that is applicable to all NMR tissues; the NMR pan-tissue clock (**Figure 1**).

**Figure 1:**
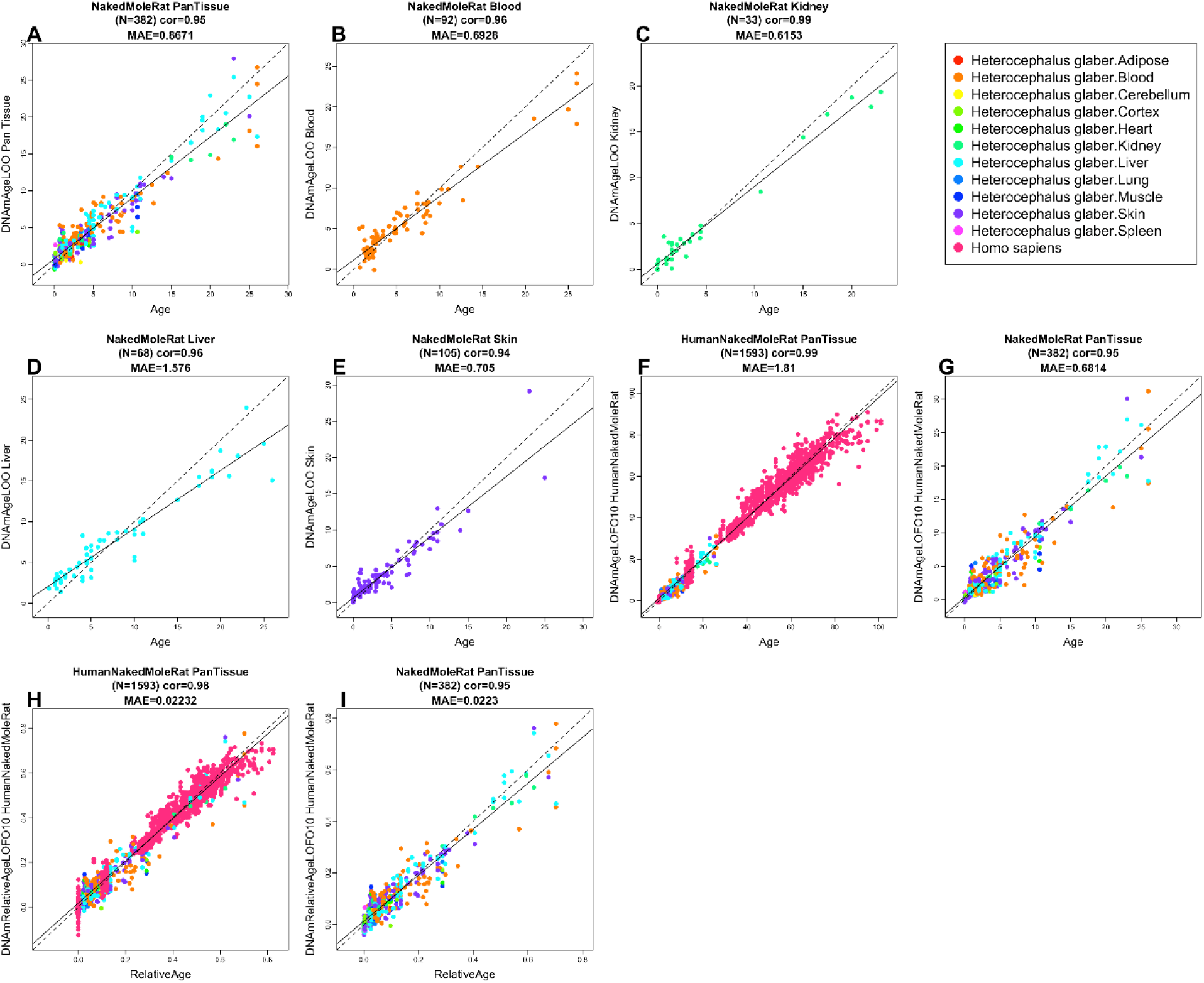
Cross-validation study of epigenetic clocks for naked mole-rats and humans. Chronological age versus cross validation estimates of DNA methylation age (y-axis, in units of years) for different epigenetic clocks A) pan tissue epigenetic clock, B) blood clock, C) kidney clock, D) liver clock, E) skin clock, F,G) human-NMR clock for chronological age, H,I) human-NMR clock for relative age. We used two validation schemes: A-E) leave-one-out cross validation (LOO) and F-I) ten-fold cross validation balanced for species (LOFO10). F,G) human-naked mole-rat clock for chronological age applied to F) both species, and G) NMR only. H,I) Human-NMR clock for relative age applied to H) both species, and I) NMR only. Dots are colored by F,H) species (red=human) or G,I) NMR tissue type. G,I) Same as panels F,H, respectively, but restricted to NMR samples. Each panel reports the sample size, correlation coefficient, median absolute error (MAE).

Pan-tissue, blood, kidney, liver and skin clocks generated very strong age correlations (R>=0.94, Figure 1A-E). A detailed analysis of the performance of the pantissue clock with different tissues is presented in **Supplementary Figure S2**. Tissues with large numbers of samples (blood, kidney, liver and skin) showed excellent correlation between DNAm age and chronological age. Overall, these NMR clocks are highly accurate estimators of chronological age that the research community can readily employ in their investigations into ageing.

As the aim of studying NMR ageing is to ultimately understand human ageing, it would be beneficial if age measurements of NMR can be readily translated to human age. Towards this end, we developed two dual-species clocks (human-NMR). These clocks were derived by employing the elastic net regression approach as described above, but on the training data set that consisted of DNA methylation profiles of both human and NMR tissues, that were all profiled with the mammalian methylation array.

The output of one of the human-NMR clocks reports age in years, while the other reports relative age, which is the ratio of chronological age of an individual to the maximum recorded lifespan of its species, and assumes values between 0 and 1. This circumvents the issue that arises when age comparisons are made, in years, between members of these two species, which have very different lifespans. Relative age on the other hand, is more meaningful as it indicates the life stage of an individual in respect to its potential maximum lifespan. The resulting human-NMR clock for chronological age (in years) was highly accurate whether both species were analyzed together (R=0.99, Figure 1F) or when the analysis was restricted to NMR (R=0.95, Figure 1G). Equally strong age correlations were obtained with the human-NMR clock for *relative age* regardless of whether the analysis was carried out with both species (R=0.98, Figure 1H) or only NMRs (R=0.95, Figure 1I). These results suggest that a subset of CpGs in both human and NMR undergo similar methylation changes during aging, allowing the derivation of highly accurate dual-species age-estimators. These clocks will be useful for readily translating discoveries made in NMR to humans. The successful generation of these epigenetic clocks implies that NMRs, despite being deemed as “non-ageing”, undergo epigenetic ageing, just like other members of the mammalian class.

### Characteristics of age-related CpGs

To uncover epigenetic features of the NMR genome that are associated with age, we first examined the location of age-related CpGs in its genome. These were found to be distributed across all genomic structures, with some CpGs increasing and other decreasing methylation with age (**Figure 3C**). Importantly, an overwhelming proportion of age-related CpGs that are located in promoters and 5’UTRs, become increasingly methylated with age. This parallels the observed hypermethylation of CpG islands with age (**Figure 3D**). As CpG islands control the activities of many promoters, their methylation suggests the potential participation of their corresponding proximal genes in the process of ageing. It is acknowledged that methylation of non-island CpGs can also influence gene expression. To identify CpGs with methylation changes that are most related to ageing, we performed an epigenome-wide association study (EWAS), where we correlated age with each of the 27,917 CpGs on HorvathMammalMethylChimammalian methylation array that mapped to specific loci in the *Heterocephalus_glaber_female.HetGla_female_1.0.100* genome assembly. In total, these CpGs are proximal to 4,634 genes in the NMR genome. The resulting EWAS appears to show strong tissue-specificity with regards to age-associated CpG methylation changes. In other words, there were very few common age-related CpGs identified between the different tissues (**Supplementary Figure S4**). This poor conservation and the differences in p-values between tissue types, however, may be due to the limited sample size of some tissues. This notion is consistent with the observation that three tissue types with larger sample size (blood, n=92; liver, n=68; and skin, n=105) showed more consistent aging effects on DNA methylation (**Figure 3A, B** and **Supplementary Figure S4**).

**Figure 2.**
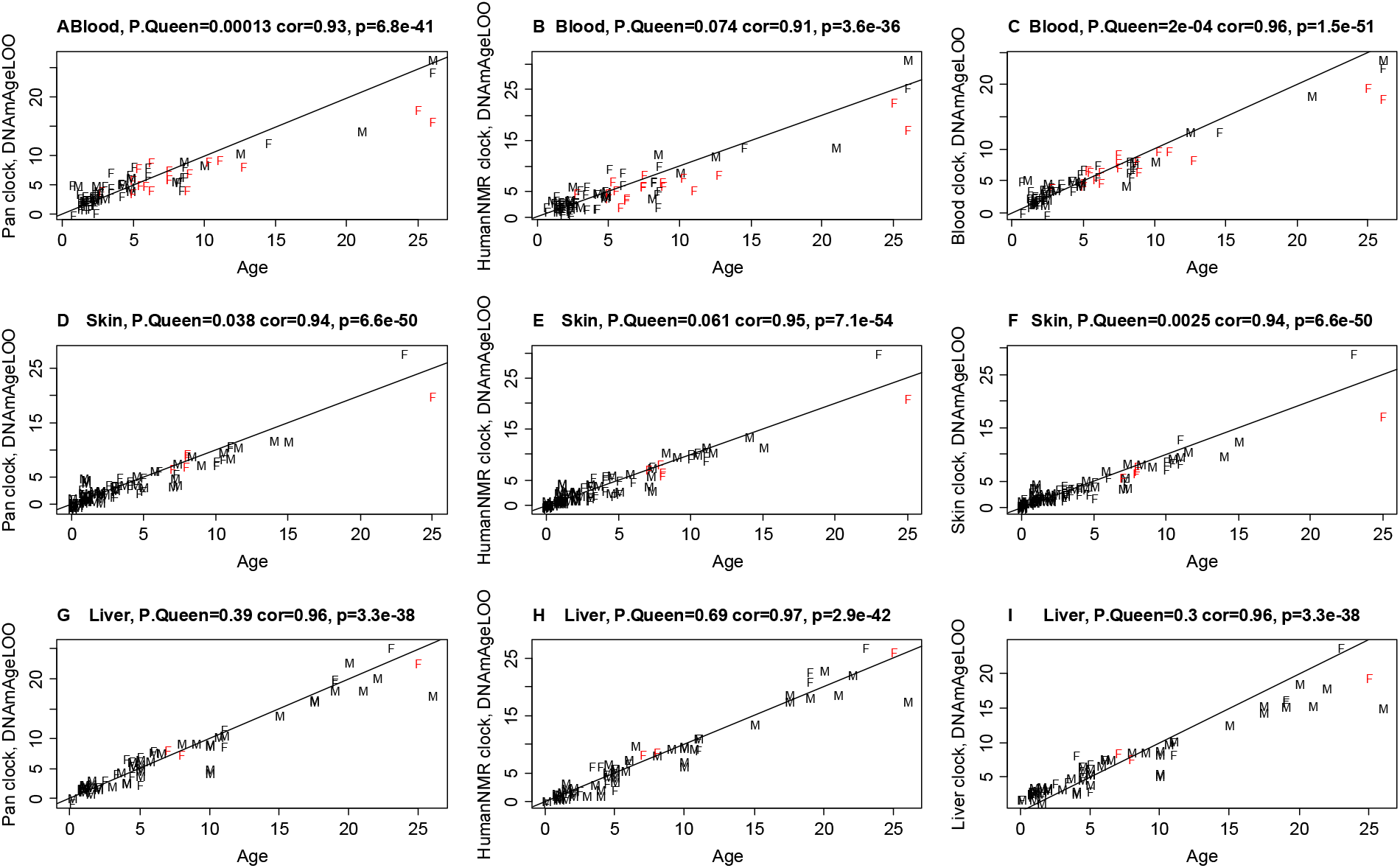
Leave-one-out estimates of DNAm age (y-axis) colored by queen status. All axes are in units of years. x-axis reports the chronological age at the time of the sample collection. Dots are labeled by sex (Female vs Male) and colored by queen status: red=queen, black=“not a queen or unknown”. Rows correspond to tissues: (A,B,C): blood, D-F) Skin, G-I (liver). Columns correspond to different clocks: first column (A,D,G) reports results for the NMR pan tissue clock. Second column (B,E,H) reports results for the human-NMR clock of chronological age. Third column reports tissue specific clocks C) NMR blood clock, F) skin clock, I) liver clock. P.Queen denotes the p-value for a differential slope analysis comparing aging patterns in queens (red dots) to the remaining samples (black dots). This p-value was calculated from a linear regression model whose dependent variable (DNAmAge) was regressed on age, queen status, and their interaction effect. P.Queen was calculated using the Wald test statistic for the interaction effect.

**Figure 3.**
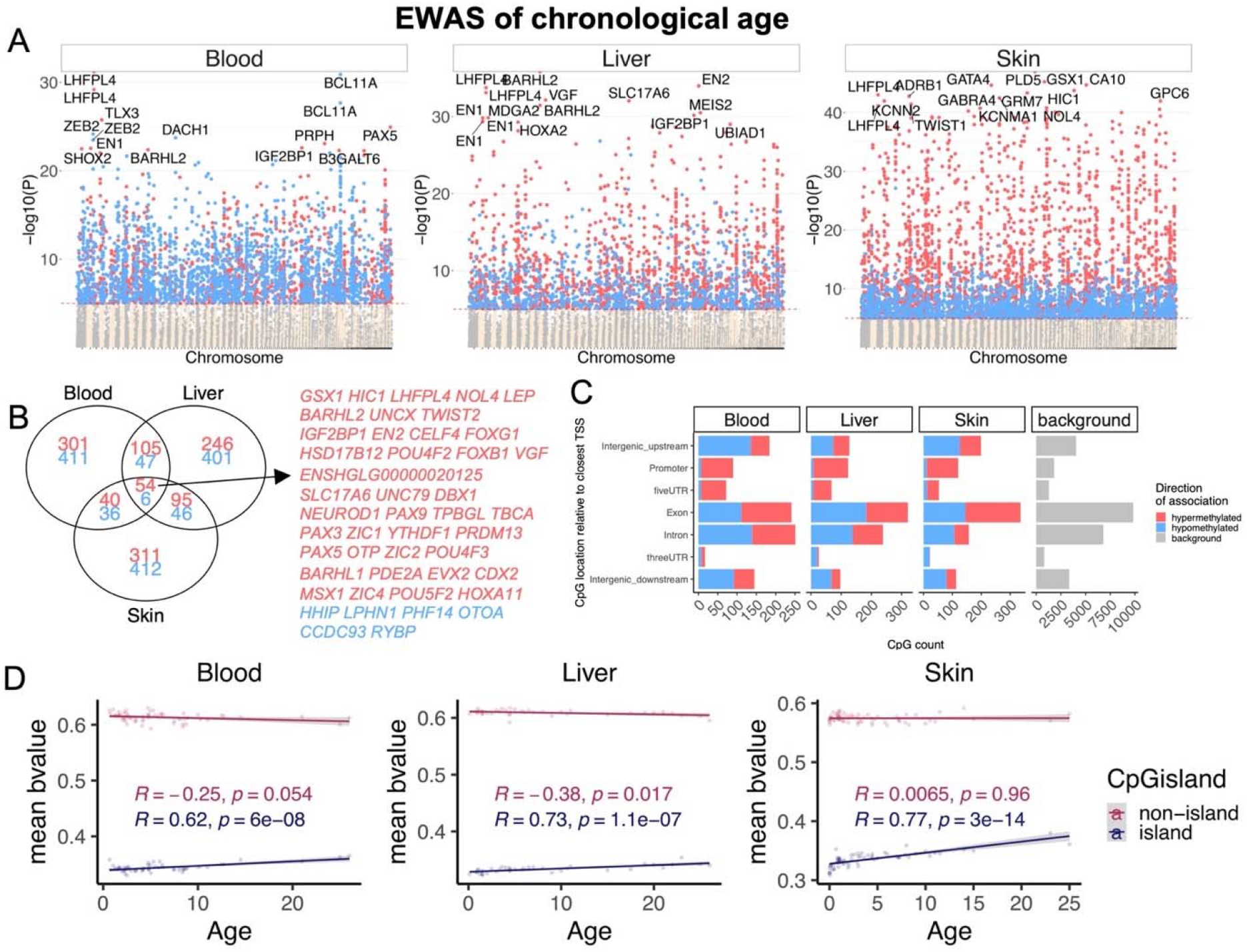
Epigenome-wide association (EWAS) of chronological age in naked mole-rat tissues. EWAS of age in blood (n=92), liver (n=68), and skin (n=105) of NMR. The results from tissues with small sample sizes (cerebral cortex, heart, kidney, lung, muscle) are in Supplementary Figure S5. A) Manhattan plots of the EWAS of chronological age. The coordinates are estimated based on the alignment of Mammalian array probes to HetGla_female_1.0.100 genome assembly. The direction of associations with p<10^−5^ (red dotted line) is highlighted by red (hypermethylated) and blue (hypomethylated) colors. Top 15 CpGs was labeled by the neighboring genes. B) Venn diagram of the overlap in up to top 1000 (500 in each direction) significant CpGs in each tissue. C) Location of top CpGs in each tissue relative to the closest transcriptional start site. The grey color in the last panel represents the location of 27917 mammalian BeadChip array probes mapped to HetGla_female_1.0.100 genome. D) Scatter plot of DNA methylation aging in CpGs located in island or non-island CpGs in NMR tissues. The plots show the average of DNAm beta values per CpG island status for each animal.

To identify top genomic loci that exhibited age-related methylation changes in each tissue, a threshold p-value < 10^−5^, was adopted. Genes that are proximal to the top scoring age-related CpGs in each tissue are as follows: blood, *LHFPL4* (CpG is located in an exon, z = 13.4); liver, *BARHL2* upstream (z = 15.3); skin, *PLD5* exon (z = 17.5). To identify common age-related CpGs across different tissues, an *upset* plot analysis was carried out. An upset plot is a generalization of Venn diagrams to more sets. We identified CpGs with conserved age-related methylation change in at least five NMR tissue types (**Supplementary Figure S5**). Genes that are proximal to these CpGs include *LHFPL4, BARHL2, SLC5A11,* and *VGF.* We are mindful of the fact that the numbers of different tissues in our analysis are different and differences in statistical power may influence the results. Hence, for a more robust identification of shared age-related CpGs, we limited this analysis to the three tissues (blood, skin and liver) for which there were more than 50 samples. This identified 60 common loci in the NMR genome that have consistent age correlations in the 3 tissues (**Figure 3B**).

To gain additional insights into age-related biological pathways we performed enrichment analyses. potential age-related genes in individual tissues. Enrichment analysis of genes adjacent to age-related CpGs in tissue types with sufficiently large sample size and also the shared 61 CpGs, revealed the potential involvement of several canonical pathways that are known to be related to the biology of ageing. These include regulation of telomerase, FOXA1-related pathway, mitochondrial function, and Wnt signaling. Other pathways highlighted were mostly related to regulation of transcription in general. This is consistent with the results of gene ontology analysis in which DNA-binding featured prominently amongst other processes (**Supplementary Figure S6**. Furthermore, the genes proximal to hypermethylated age-related CpGs are locations of H3K27Me3 marks, polycomb protein EED binding sites, and PRC2 targets. It is acknowledged that CpG methylation can also influence expression of more distal genes. The absence of any means to identify these genes, however, precludes their analysis.

### Epigenetic analysis of queen status

NMRs, together with Damaraland mole-rats, have been proposed to be eusocial mammals. Eusocial species have very organized lifestyles and clear division of labor and roles, with the queens (the dominant breeding female in the colony) being the most distinct members of the social group. Given the genetic similarity between queens and non-breeders, it is highly likely that eusocial characteristics could be determined by, or correlated with, epigenetic differences. Hence, we investigated the potential relationship between queen status and DNA methylation. As we had blood DNA from 18 queens, we carried out this analysis primarily with blood DNA, and a subset of analyses also included skin, for which there were only 4 queen samples. We successfully identified 41 CpGs in blood and 232 in skin that were differentially methylated between queen and non-breeding females at a significance of α=10^−4^ (**Figure 4A**). Many of them in regions that encode transcription factors that are involved in neural/brain processes, which may possibly contribute to differential behavior between queens and non-breeders. These sets of genes will be of interest for researchers interested in NMR behavior differences between queen and non-breeders.

**Figure 4.**
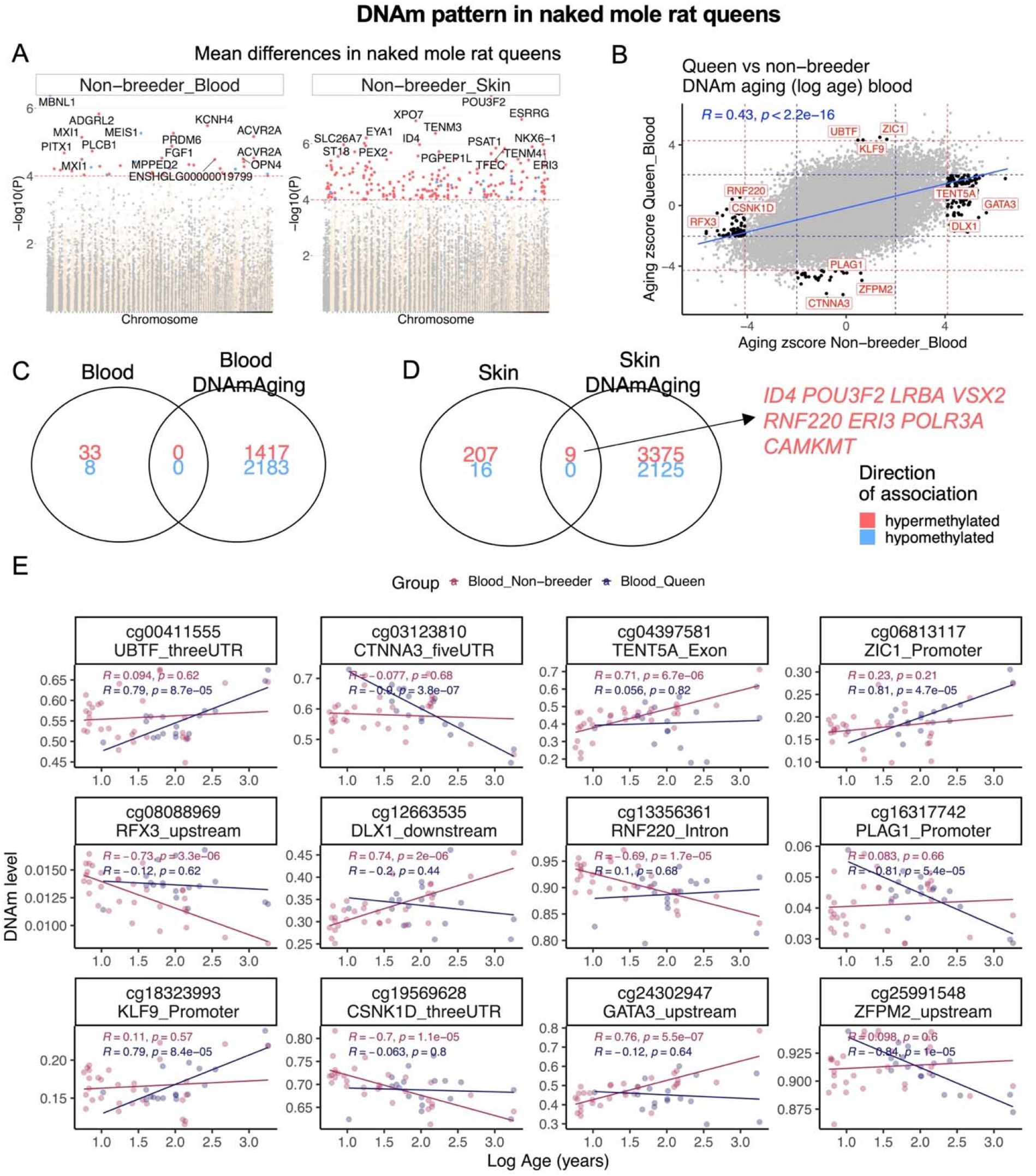
Queen status modestly alters DNA methylation pattern in NMR. A) Manhattan plot of the EWAS of queen status (Queen vs non-breeder females) in blood and skin of NMR. Sample sizes: non-breeding female blood, 32; non-breeding female skin, 8; Queen blood, 18; Queen skin, 4. The EWAS analysis used chronological age as covariate in order to control this potential confounder. The red line in the Manhattan plot corresponds to a significance level of p =1e-4. B) Sector plot of the CpGs with distinct DNA methylation aging in the blood of queen and non-breeder females. We carried out a log transformation (natural log) of age since the age distribution of queens was highly skewed. The black dots indicate the CpGs that change in one (p<10^−4^, red dotted line) and do not change (p>0.05, blue dotted line) in the other analysis. Skin tissue was excluded from this analysis due to limited sample size for a stratified EWAS of age. Venn diagrams of the overlap between queen effects in blood (C), skin (D), and significant CpGs with DNAm aging in each tissue. E) DNAm aging effects in blood samples for selected CpGs that differ between queen and non-breeding females.

### Slower epigenetic aging rates in queens

With the highly accurate epigenetic clocks in hand, we were well-placed to ascertain whether queens (i.e. reproducing females) differ from female and male nonbreeders with regards to epigenetic aging rates. When tested, queens were indeed found to age more slowly than non-breeders, as measured by leave-one-out estimates of DNAm age for blood and skin (**Figure 2**). This was particularly clear from analysis of blood samples, for which there were more queen samples than for other tissues (p=0.00013 with pan-tissue clock Figure 2A, p=0.074 with human-NMR clock Figure 2B, p=0.0002 with blood clock Figure 2C). There was nominally significant evidence of slower aging rates in skin samples from 4 queens, the oldest of which showed substantially younger epigenetic age as predicted by the skin clock (**Figure 2**) (p=0.038 for the pan-tissue clock Figure 2D, p=0.061 for the human-NMR clock Figure 2E, p=0.0025 for the skin clock). Age estimates of other tissues such as liver were not significant and inconclusive due to even smaller sample size (Figure 2G-I). The abovementioned analysis has a statistical limitation: both queens and non-queens were used to arrive at the leave-one-out estimates of DNAm age. Since the training sets contained queens, the resulting clock may inadvertently condition out the slower aging effects of queens. To address this concern, we employed the second analysis strategy. We used male samples as training data and female data as test data. Thus, we first developed epigenetic clock for blood and skin from *male* NMRs (training set). We then used this male clock to evaluate female blood and skin samples (female test set). Queens were once again observed to exhibit slower epigenetic aging, according to visual inspection (**Supplementary Figure S3C**) and by multivariate linear regression model analyses of female samples (Wald test p=0.015). Since we were concerned that samples from older queens may drive the observed association, we repeated the analysis in animals who were younger than 15. Again, a multivariate regression analysis revealed that queens aged more slowly than female non-queens (p=0.031).

Age-related CpG methylation changes in queen blood were found to correlate well (R=0.43) with those of non-breeder females (**Figure 4B**), indicating broad conservation of epigenetic ageing mechanism between them, despite the differences in aging rate. However, we did identify a total of 237 age-related CpGs with unique methylation patterns in either of these groups (**Figure 4B, E**). It is possible that such mutually exclusive age-related CpGs may contribute to the different rate of epigenetic ageing between queens and non-breeders. This remains to be empirically tested, although enrichment analysis of these 237 unique age-related CpGs highlighted the involvement of developmental pathways, particularly those of sensory perception. Interestingly, the LHX3 transcription factor motif was also identified by the enrichment analysis (**Supplementary Figure S7)**. LHX3 transcription factor controls pituitary development and may be responsible for maintaining both reproductive status and aging characteristics of the animals. In skin, we identified 9 CpGs that were associated with queen status as well as age (**Figure 4D**). These CpGs could potentially contribute to epigenetic age differences in the skin of these two groups of NMR, although it is necessary to be mindful of the limited number of skin samples and the fact that we did not identify in blood, any age-related CpGs with baseline methylation that differed between queen and non-breeders (**Figure 4C**). Collectively, the various analyses above have engendered an interesting list of target genes to be tested for their potential contribution in regulating NMR ageing, queen status and behavior.

### Age effects in primates, mouse and naked mole-rat tissues

As mentioned above, a major unique feature of NMR is their outstanding longevity compared to other small rodents, such as the mouse. Humans are another species with extraordinarily long lifespan. As such NMR and humans are joint outliers with regards to expected lifespan. Hence, it would be interesting to identify age-related CpGs that are shared between NMRs and humans, but not with mice, which are phylogenetically closer to the NMR. The identification of genes that are proximal to these CpGs may enlighten us about the molecular and cellular processes that underpin the longevity of NMR and humans. Towards this end, we compared age-associated CpGs in three tissues (liver, skin and blood), for which there were sufficiently large number of samples in all three species (Table 1). We did not, however, utilize human liver data because the human liver samples in our possession were from older adults while many NMR livers were from young animals. This would introduce a bias that would have affected the results. As such we used liver data from another primate: the vervet monkey, whose liver samples were profiled uniformly across the entire lifespan^36^.

In general, we observed that age-related CpGs of NMR overlapped moderately with those of mice and primates (human and vervet monkey, **Supplementary Figure S8–S12**). In particular, age-related CpGs of NMR blood correlated weakly (r=0.2) with those of mouse and human (**Supplementary Figure S8C, D**). There were however, better correlations between age-related CpGs of NMR skin (~ r=0.4) and liver (~ r=0.5) with those of mouse and primates. Interestingly, age-related CpGs of NMR correlated much better with those of vervet monkey (r=0.6) than with those of mouse (r=0.4), despite the closer phylogenetic relationship of the latter to NMR (**Supplementary Figure S8C, D**).

To identify age-related CpGs associated with species longevity we focused on those that change with age similarly between NMR and primates, but differentially in the mouse. In total, subsets of 171 such CpGs were detected in liver, 50 in blood, and 116 in skin (**Supplementary Figure S13**). Identification of genes that are proximal to these cytosines revealed several that are of particular interest. Some of these that were identified in at least two of the three tissues include; hypermethylation of the *IGF2BP1* promoter, *PAX2* upstream, SLITRK1 promoter, and *B3GALT6* exon. Strikingly, these regions were highly enriched in pathways related to lethality and abnormal survival in mouse phenotype database (**Supplementary Figure S14**). In general, a prominent feature of age-related CpGs that are shared between NMRs and primates but not the mouse, is their proximity to genes that encode proteins with homeobox domains, particularly the NKL and HOXL family of proteins (**Table S14**). These proteins are primarily transcription factors that regulate expression of developmental genes. As such it follows that their perturbations in mice were reportedly associated with lethality during fetal growth, post-natal lethality, abnormal survival, decreased body size, premature ageing and death, nervous system abnormality etc. (**Table S14**). Besides development, genes encoding metabolic proteins were also found to be proximal to age-associated CpGs shared between NMRs and primates but not the mouse. Collectively, these results suggest that differences in development and metabolism may in part, jointly underlie the longevity of NMR and primates (especially humans), which is longer than expected from their body-size. These identified genes (**Table S14**) are valuable targets for experimental perturbations to interrogate life expectancy or aging-related pathologies.

## DISCUSSION

This study describes seven epigenetic clocks for NMRs, of which five are specific to naked mole-rats (for different tissue types) and two are dual-species human/naked mole-rat clocks that are applicable to humans as well. The human-naked mole-rat clocks for chronological and relative age demonstrate the feasibility of building epigenetic clocks for different species based on a single mathematical formula. This further consolidates emerging evidence that epigenetic aging mechanisms are conserved, at least between members of the mammalian class.

The first step towards developing these clocks was the removal of the species barrier that precluded the use of EPIC arrays to profile non-human species. In response, we developed the mammalian methylation array that profiles 36k CpGs with flanking DNA sequences that are highly conserved across numerous mammalian species. This allowed us to generate DNA methylation profiles from different species that can be directly analysed either individually or collectively, as they were all derived using the same DNA methylation measurement platform. This is a technical step-change from previous NMR studies, which utilized sequence-based measurement platforms^18,37^, which do not lend themselves to direct analysis and comparison with other species.

At face value, two clearly established facts appear to contradict each other. On a phenotypic level, the NMR appears to evade aging. Yet our study clearly detects significant age-related changes in DNA methylation levels across the entire lifespan of the animal, even in relatively young ones. Indeed, the age effects are so pronounced that they allowed the development of highly accurate age estimators based on DNA methylation levels that apply to the entire lifespan of the NMR. These two seemingly contradictory results can be interpreted in different ways. First, they could imply that age-related DNA methylation changes do not matter since they do not appear to correlate with any adverse functional consequences in NMR. This perspective however is difficult to reconcile with the fact that accelerated epigenetic ageing has been correlated to a very wide range of pathologies and health conditions^38^. Alternatively, it could mean that while the NMR ages at a molecular level, as do all other mammals, it has developed compensatory mechanisms that counteract the consequence of these epigenetic changes. The NMR age-related CpGs that we identified, and the availability of epigenetic clocks, are new and valuable resources that will help bring resolution to this question.

Further clues to NMR ageing were also revealed from the three-way comparison of age-related CpGs between NMRs, primates and mouse. Although primates and NMRs are phylogenetically more distantly related than NMRs and the mouse, these relationships are not similarly manifested when it comes to longevity. Indeed, NMRs and humans are more akin to each other as they are both outliers with regards to lifespan expected from their adult size. Here, the three-way comparison revealed that the unusually long lifespans of NMR and primates may lie in the co-regulation of developmental and metabolic processes. This is a fascinating perspective as these two distinct biological processes have been studied, investigated and considered mostly in isolation of each other. The observations reported here indicate that a comprehensive understanding of ageing and longevity may require the integration of our current understanding and future investigation of these two biological processes. Conversely, similarly regulated developmental genes between NMR and human may reflect neotenic features characteristic of these two species^39^ Neoteny is defined as retention of juvenile features into adulthood. A shift towards longer development and retention of youthful tissue repair can lead to longevity.

With regards to NMR lifestyle, we identified a large number of CpGs in blood and skin that are differently methylated between queen and non-breeding females. More than 99% of NMRs will never reproduce in their long lifespan due to their characteristic reproductive division of labor^40^ Indeed, the reproductive tracts of non-breeders of both sexes remain in a pre-pubertal state throughout adulthood till death (for review see^41^). However, NMRs are “totipotent” in that most, if not all, are apparently capable of becoming reproductively active should they be removed from social cues that suppress their reproduction (namely, the presence of a breeding queen). Because of this plasticity in reproductive capacity, epigenetic effects seem likely to play a role in determining queen/non-breeder status and in mediating social suppression of reproduction. Remarkably, demographic data from a large number of captive bred NMRs showed longer survival of breeders^11^. This is consistent with the observed slower epigenetic ageing of queens, which argues for biological relevance of age-related epigenetic changes in NMR, as was considered above. Further analysis of DNA methylation aging in function of social status (queen vs non-breeders) implicated the transcription factor LHX3, as a potentially important ageing rate determinant that plays important roles in both pituitary and central nervous system development^42–44^. LHX3 binds to promoters of several genes involved in pituitary development and function, including αGSU, TSHß, FSHß, PRL, gonadotropin-releasing hormone receptor, and PIT-1^45–49^, which could potentially regulate lifespan. Strikingly, alterations in pituitary development was previously shown to lead to slow aging phenotype in Ames and Snell dwarf mice^50,51^.

In summary, we have developed robust epigenetic clocks for NMRs. These clocks can be used to estimate the age of wild NMRs, and more excitingly to facilitate the studies of NMRs as a model organism for ageing, longevity and suppression of pathologies. The dual human-NMR clocks are expected to be particularly valuable as they allow for the direct translation of findings in NMRs to humans. Furthermore, our results demonstrate that NMRs age epigenetically despite displaying negligible senescence. This finding underscores that exceptionally long-lived species may evolve mechanisms to uncouple epigenetic aging from physiological decline. We have also identified genes and cellular pathways that impinge on developmental and metabolic processes that are potentially involved in NMR and human longevity. Finally, we demonstrate that NMR queens age more slowly than non-breeders and identify several genes including LHX3 transcription factor involved in pituitary development to be associated with queens’ longevity.

## EXPERIMENTAL PROCEDURES

The NMR tissue samples were provided by two different labs: 1) Vera Gorbunova and Andrei Seluanov from the University of Rochester and 2) Chris Faulkes from the Queen Mary, University of London.

### Animals from the University of Rochester

All animal experiments were approved and performed in accordance with guidelines set up by the University of Rochester Committee on Animal Resources with protocol number 2009-054 (naked mole-rat). Naked mole-rats were from the University of Rochester colonies. All animals in the colonies are microchipped and their ages are recorded. Housing conditions are described previously^52^. All tissues except for skin biopsies and blood were obtained from frozen tissue collection at the University of Rochester from healthy animals that were euthanized for other studies. Skin biopsies (2 mm punch) were collected from the backs of the animals under local anesthesia. Blood samples were collected from the tails. Genomic DNA was extracted using Qiagen DNeasy Blood and Tissue kit and quantified using Nanodrop and Qubit.als

### Study Animals from Queen Mary University

Naked mole-rats were maintained in the Biological Services Unit at Queen Mary University of London in accordance with UK Government Animal Testing and Research Guidance. The tissues used in this study were obtained from post-mortem specimens from animals free from disease in compliance with national (Home Office) and institutional procedures and guidelines. Because sample collection was from post-mortem material, additional local ethical approval was not required for this study. Tissue samples were snap frozen in liquid nitrogen following dissection and transferred for storage at −80°C.

### Human tissue samples

To build the human-naked mole-rat clock, we analyzed from methylation data from 1207 human tissue samples (adipose, blood, bone marrow, dermis, epidermis, heart, keratinocytes, fibroblasts, kidney, liver, lung, lymph node, muscle, pituitary, skin, spleen) from individuals whose ages ranged from 0 to 93 years. Tissue and organ samples were obtained from the National NeuroAIDS Tissue Consortium^53^, blood samples from the Cape Town Adolescent Antiretroviral Cohort Study^54^. Blood, skin and other primary cells were provided by Kenneth Raj^55^ Ethics approval (IRB#15-001454, IRB#16-000471, IRB#18-000315, IRB#16-002028).

### DNA methylation data

The mammalian DNA methylation arrays were profiled using a custom Infinium methylation array (HorvathMammalMethylChip40) based on 37492 CpG sites. Out of these sites, 1951 were selected based on their utility for human biomarker studies; these CpGs, which were previously implemented in human Illumina Infinium arrays (EPIC, 450K), were selected due to their relevance for estimating age, blood cell counts or the proportion of neurons in brain tissue. The remaining 35541 probes were chosen to assess cytosine DNA methylation levels in mammalian species^36^. The particular subset of species for each probe is provided in the chip manifest file can be found at Gene Expression Omnibus (GEO) at NCBI as platform GPL28271. The SeSaMe normalization method was used to define beta values for each probe^56^.

### Penalized Regression models

Details on the clocks (CpGs, genome coordinates) and R software code are provided in the Supplement. Penalized regression models were created with glmnet^57^ We investigated models produced by both “elastic net” regression (alpha=0.5). The optimal penalty parameters in all cases were determined automatically by using a 10 fold internal cross-validation (cv.glmnet) on the training set. The alpha value for the elastic net regression was set to 0.5 (midpoint between Ridge and Lasso type regression) and was not optimized for model performance.

We performed a cross-validation scheme for arriving at unbiased estimates of the accuracy of the different DNA methylation based age estimators. One type consisted of leaving out a single sample (LOOCV) from the regression, predicting an age for that sample, and iterating over all samples.

A critical step is the transformation of chronological age (the dependent variable). While no transformation was used for the pure naked mole-rat clocks, we did use a log linear transformation of age for the dual species clocks of chronological age.

### Relative age estimation

To introduce biological meaning into age estimates of NMRs and humans that have very different lifespans, as well as to overcome the inevitable skewing due to unequal distribution of data points from NMRs and humans across age ranges, relative age estimation was made using the formula: Relative age= Age/maxLifespan, where the maximum lifespan for the two species (human=122.5 years and NMR=37 years) were chosen from an updated version of the anAge data base^58^.

### Epigenome wide association studies of age

EWAS was performed in each tissue separately using the R function “standardScreeningNumericTrait” from the “WGCNA” R package^59^. Next the results were combined across tissues using Stouffer’s meta-analysis method. The analysis was done using the genomic region of enrichment annotation tool^60^. The gene level enrichment was done using GREAT analysis and human Hg19 background^60^.

### Gene ontology enrichment analysis

The analysis was done using the genomic region of enrichment annotation tool^47^ The gene level enrichment was done using GREAT analysis^47^ and human Hg19 background. The background probes were limited to 16,801 probes that were mapped to the same gene in the NMR genome.

## ACKNOWLEDGEMENTS

This work was supported by the Paul G. Allen Frontiers Group (SH).

## AUTHOR CONTRIBUTIONS

Contribution of DNA samples and phenotypic data: Vera Gorbunova, Chris Faulkes, Andrei Seluanov, Nicholas Macoretta, Julia Ablaeva, Masaki Takasugi, Yang Zhao, Elena Rydkina, Zhihui Zhang, Stephan Emmrich. Statistical analysis: AH, JAZ, SH, CZL, JZ. Drafting and editing the article: SH, AH, VG, CF, KR. All authors edited the article. SH conceived of the study.

## DATA AVAILABILITY

All codes and data will be posted on Gene Expression Omnibus upon acceptance.

## Supplementary Figures

**Supplementary Figure S1.**
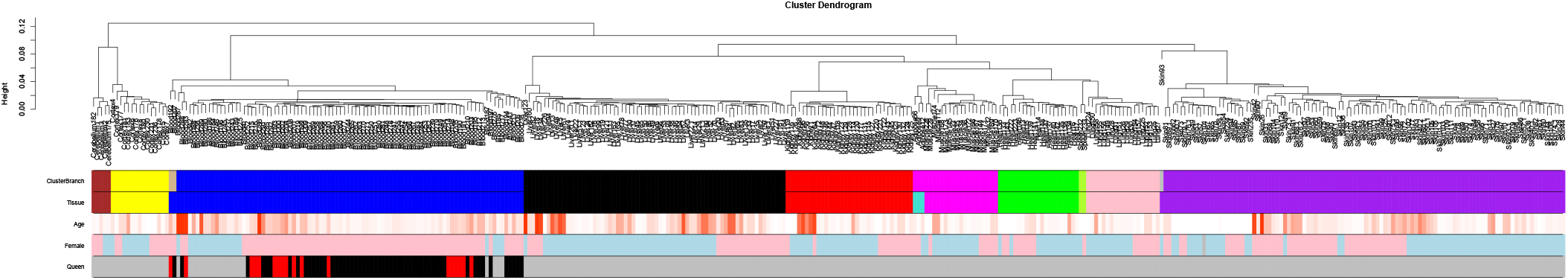
Unsupervised hierarchical clustering of NMR tissues. Average linkage hierarchical clustering based on the interarray correlation coefficient (Pearson correlation). Contrasting the first color band (based on cluster branches) with the second color band shows that the arrays cluster by tissue type.

**Supplementary Figure S2.**
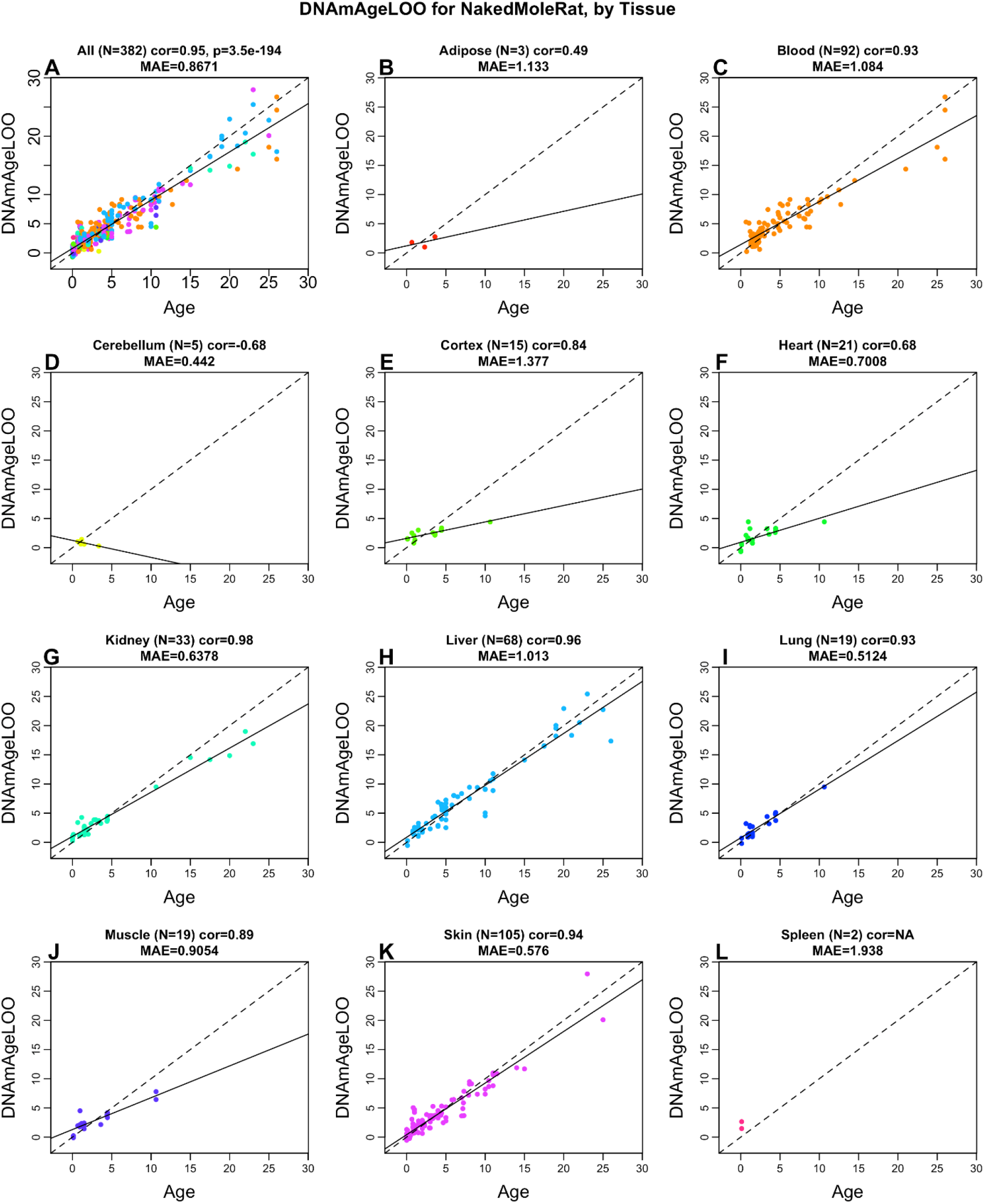
Pan tissue clock for naked mole-rat applied to individual tissues. A) All tissues combined. B) Adipose, C) Blood, D) Cerebellum, E) Cerebral cortex, F) Heart, G) Kidney, H) Liver, I) Lung, J) Muscle, K) Skin, L) Spleen. Each panel reports the leave-one-out estimate of age (y-axis) versus chronological age (in years). Each title reports the sample size, Pearson correlation coefficient, and median absolute error. The solid line results from a linear regression model. Note that panels B,D,L involve very few samples.

**Supplementary Figure S3.**
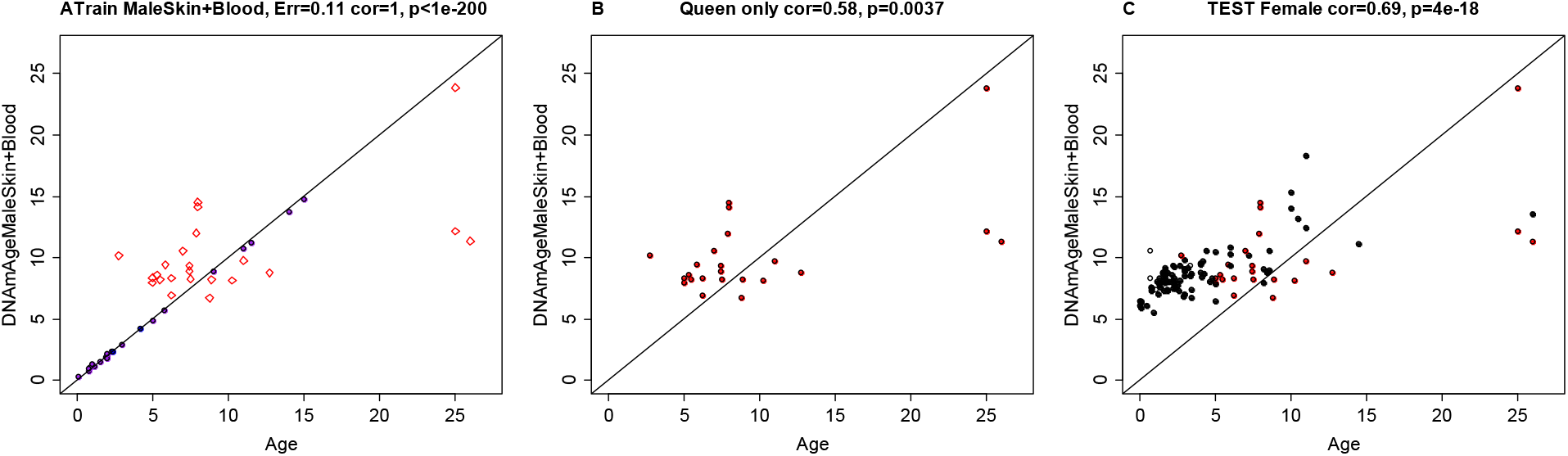
Epigenetic clock analysis of queen status in female skin and blood samples. A-C) Epigenetic clock trained on male skin and blood samples. A) Training set. The black samples correspond to the training data, i.e. blood and skin samples from male NMRs. Red diamonds correspond to part of the test data: blood and skin samples from queens. The latter data are also shown in a separate plot: panel B. C) Entire test data set. female skin and blood samples from queens and female laborers. Dots are colored by queen status: red=queen, black=non reproducing females, white=queen status is unknown. A multivariate linear regression model in female samples reveals that queen status is associated with lower epigenetic aging (p=0.0148) according to the male NMR clock.

**Supplementary Figure S4.**
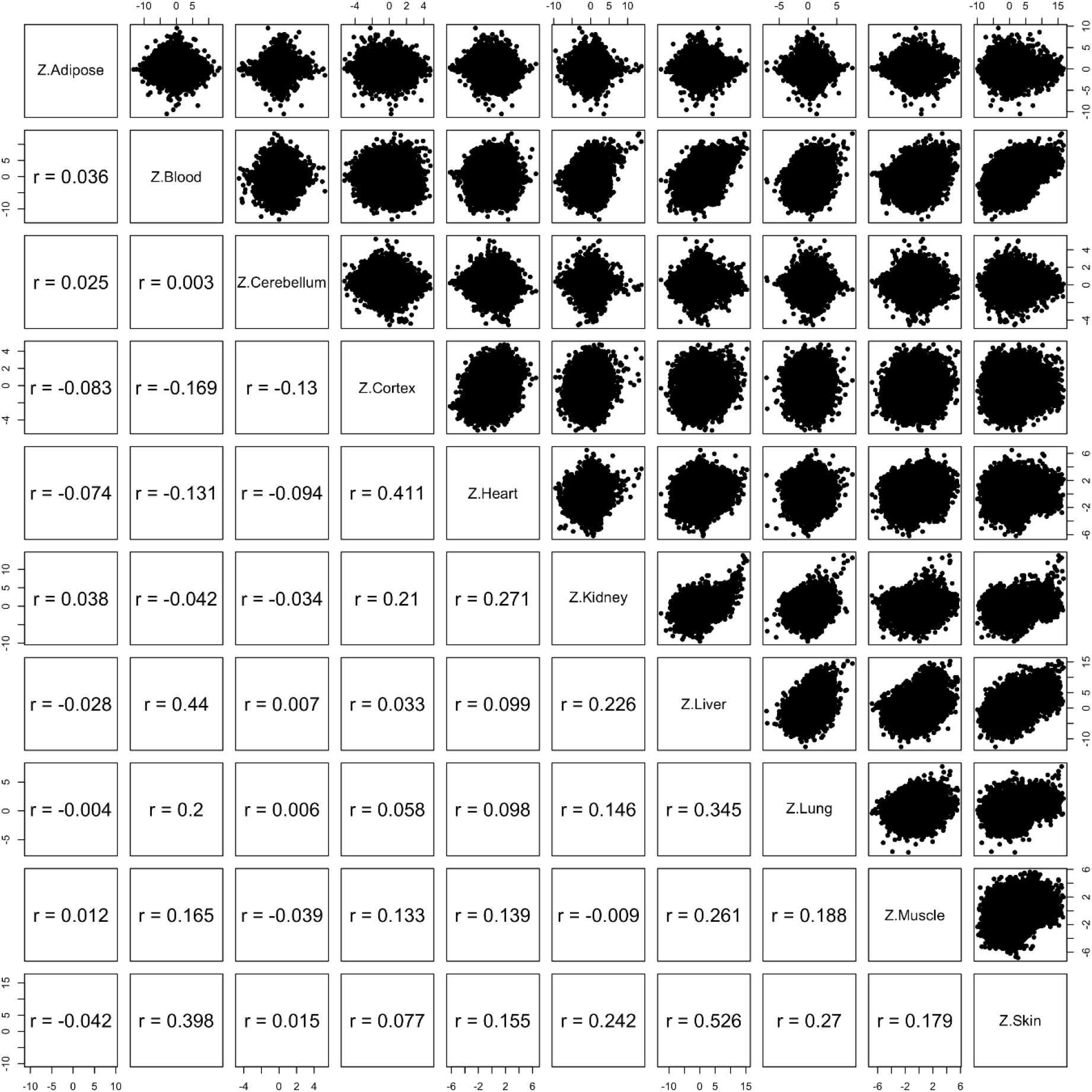
Epigenome wide association study of correlation in different tissues. Each dot corresponds to a CpG. Z statistics for a correlation test of age in the respective tissues.

**Supplementary Figure S5.**
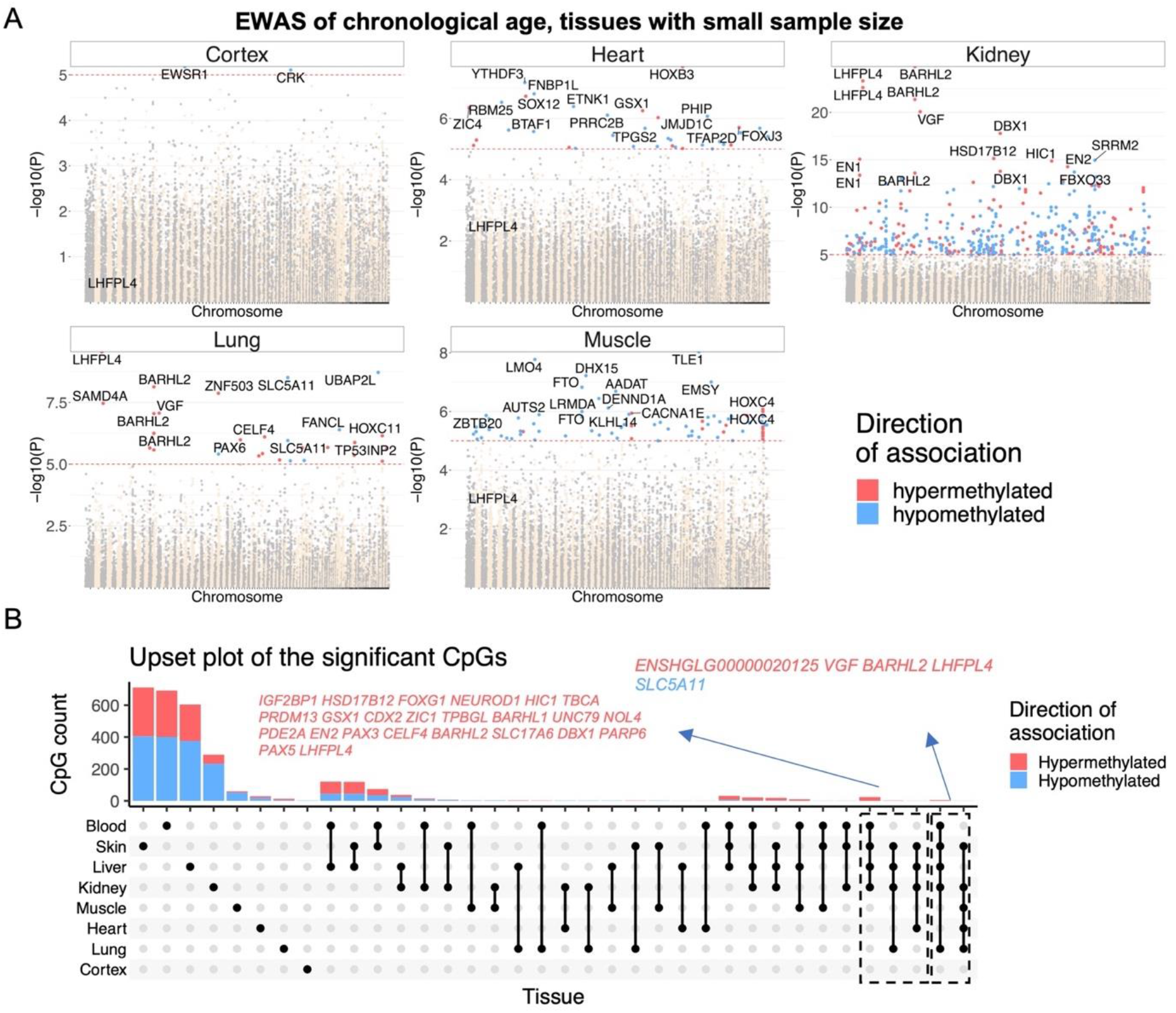
DNA methylation aging marks in NMR tissues. A) Manhattan plots of the EWAS of chronological age in tissues with small sample size (cortex, 15; heart, 21; kidney, 33; lung, 19; muscle, 19). The coordinates are estimated based on the alignment of Mammalian array probes to HetGla_female_1.0.100 genome assembly. The direction of associations with p < 10^−5^ (red dotted line) is highlighted by red (hypermethylated) and blue (hypomethylated) colors. Top 15 CpGs was labeled by the neighboring genes. B) Venn diagram of the overlap in significant CpGs in each tissue. B) Upset plot representing the overlap of aging-associated CpGs based on individual tissues.

**Supplementary Figure S6.**
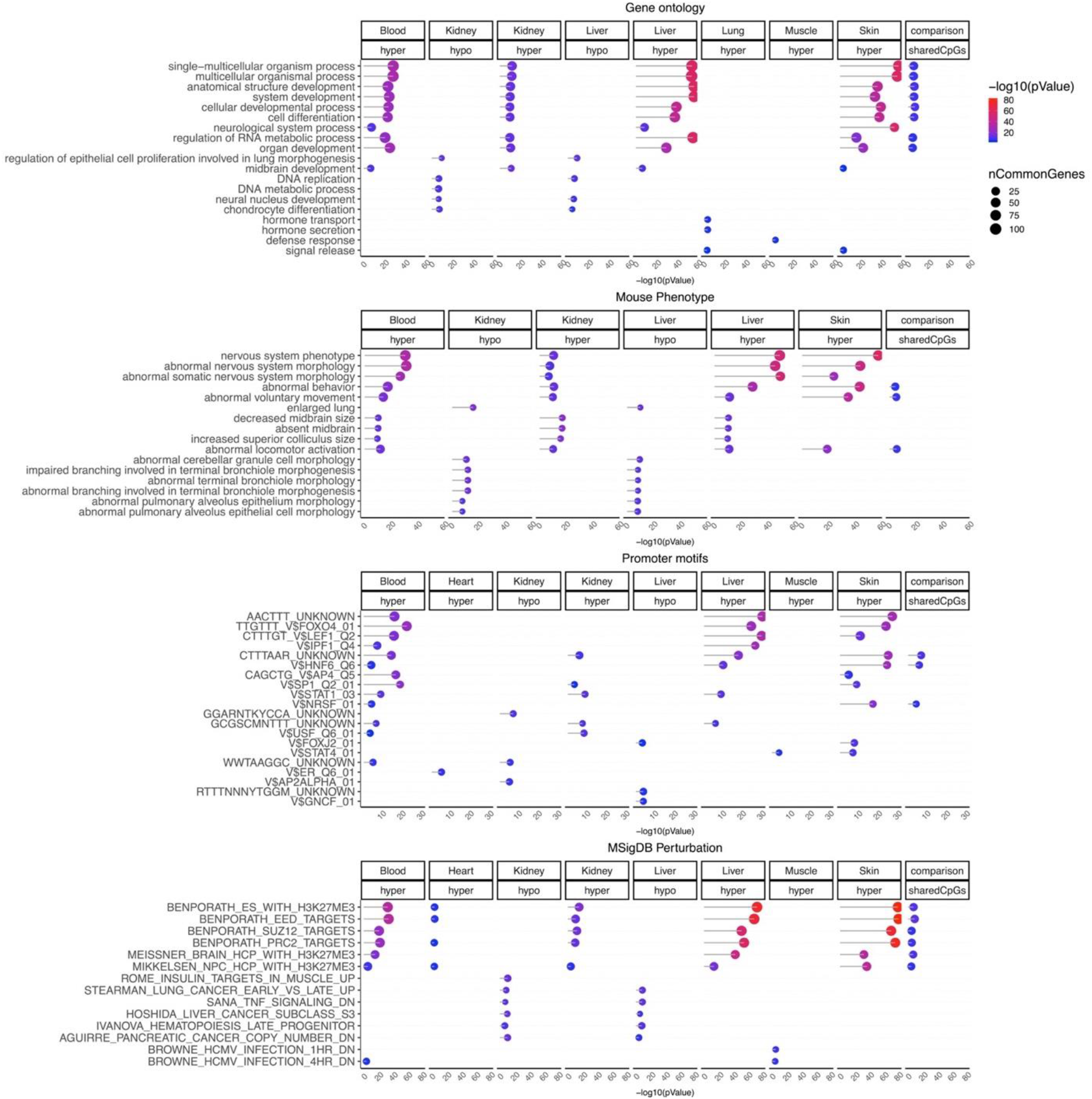
Gene set enrichment analysis of DNA methylation aging in different NMR tissues. The analysis was done using the genomic region of enrichment annotation tool^60^ The gene level enrichment was done using GREAT analysis^60^ and human Hg19 background. The background probes were limited to 14,764 probes that were mapped to the same gene in the NMR, and human. The top three enriched datasets from each category (Canonical pathways, diseases, gene ontology, human and mouse phenotypes, promoter motifs, and molecular signatures database) were selected and further filtered for significance at p < 10^−8^.

**Supplementary Figure S7.**
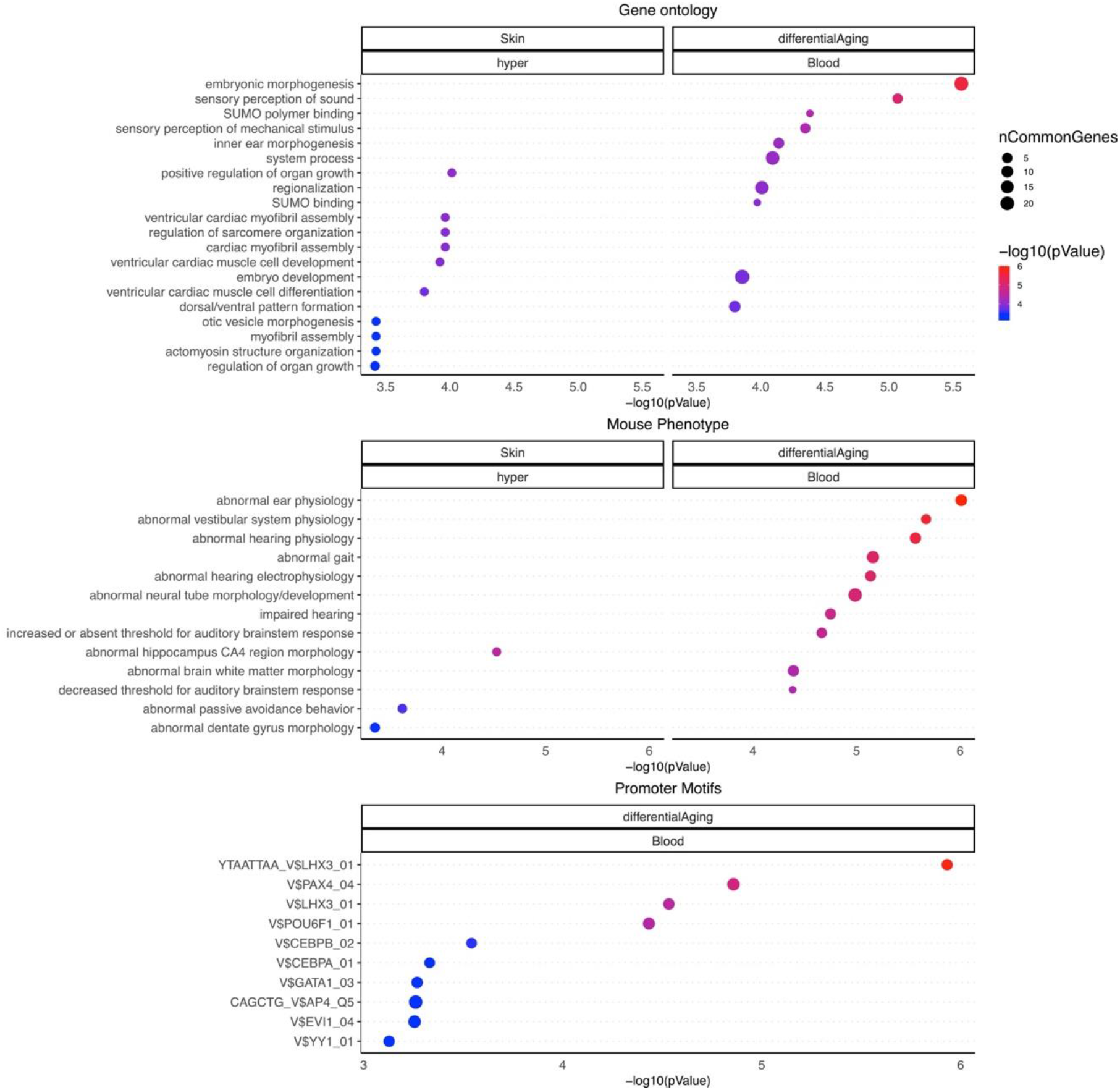
Gene set enrichment analysis of NMR female queen status basal effects and CpGs with distinct DNA methylation aging. The analysis was done using the genomic region of enrichment annotation tool^60^. Inputs: Skin/hyper, the CpGs with mean (basal) difference between queen and non-breeders at all ages; differntialAging/Blood, the CpGs with different aging pattern in blood of queen and nonbreeder females. The gene level enrichment was done using GREAT analysis^60^ and human Hg19 background. The background probes were limited to 14,764 probes that were mapped to the same gene in the NMR and human. The top 10 enriched datasets from each category (gene ontology, human and mouse phenotypes, and upstream regulators) were selected and further filtered for significance at p < 10^−3^.

**Supplementary Figure S8.**
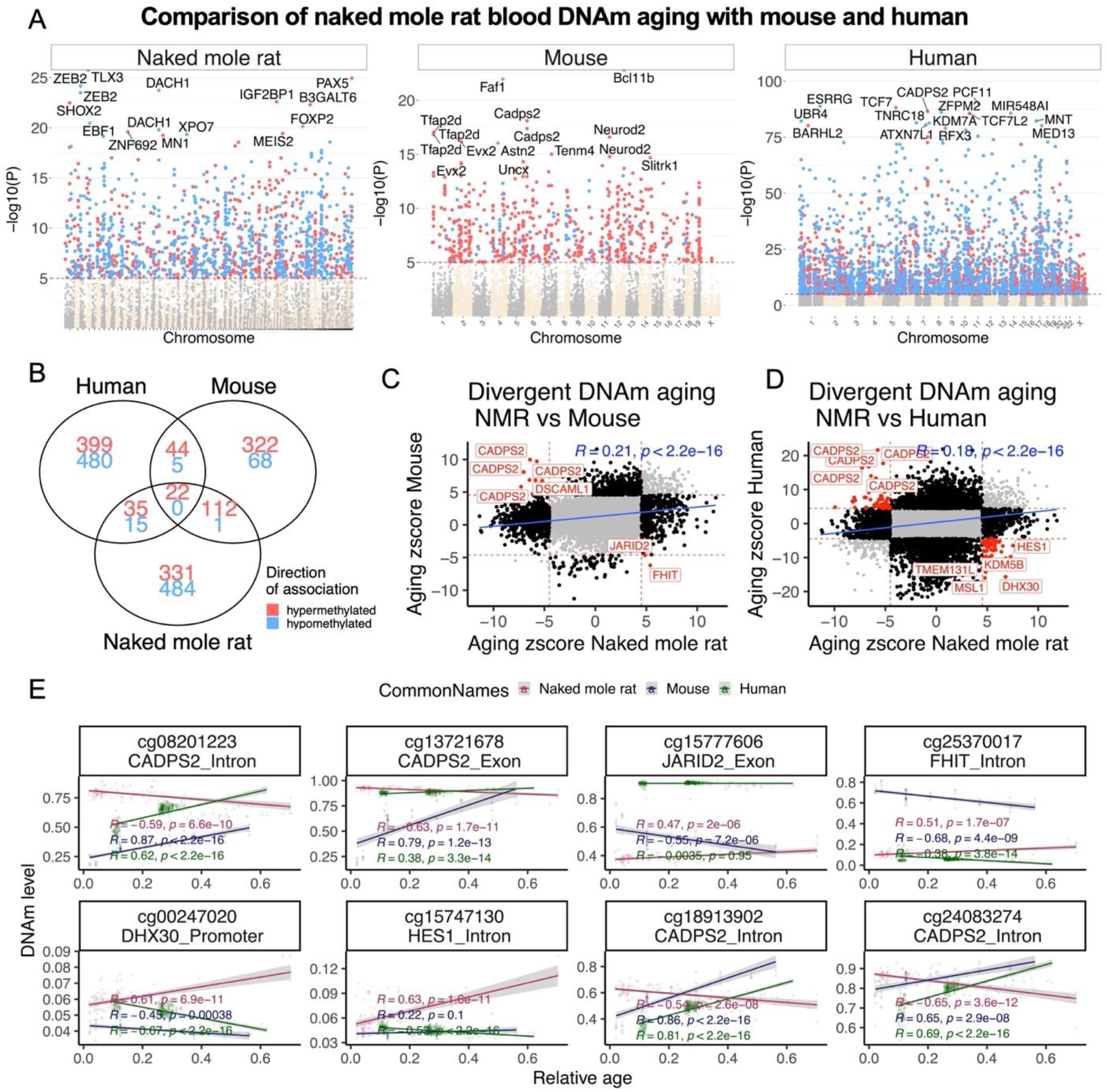
Aging effects on blood methylation data from the naked mole-rat, mouse and humans. A) Manhattan plots of DNA methylation aging loci in NMR, human and mouse. The red line in the Manhattan plot indicates p <10^−5^. The analysis was limited to 11,814 probes that aligned to orthologous genes in all three species. Sample sizes: NMR, 92; Mouse, 58; and human, 367. The red line in the Manhattan plot indicates p <10^−5^. B) Venn diagram of the overlap of significant CpGs (p < 10^−5^) in three species. Scatter plots DNA methylation aging in NMR, and mouse (C), and human (D). The highlighted CpGs are the loci that diverged in aging pattern between NMR and other species. E) DNA methylation aging in selected loci with divergent aging pattern between NMR, and other species.

**Supplementary Figure S9.**
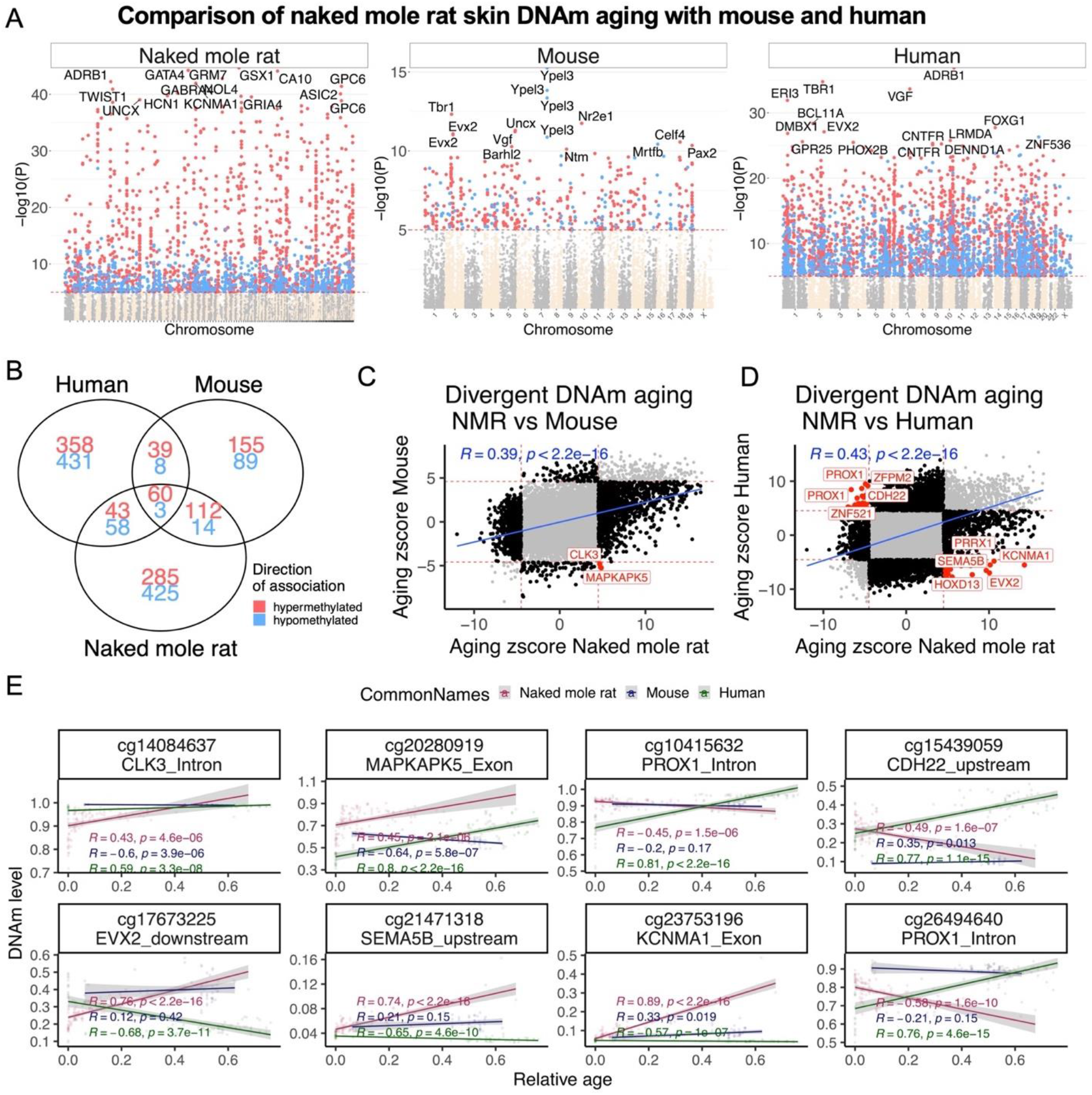
Aging effects on skin methylation data from the naked mole-rat, mouse and humans. A) Manhattan plots of DNA methylation aging loci in NMR, human and mouse. The red line in the Manhattan plot indicates p <10^−5^. The analysis was limited to 11,814 probes that aligned to orthologous genes in all three species. Sample sizes: NMR, 104; Mouse, 42; and human, 72. The red line in the Manhattan plot indicates p <10^−5^. B) Venn diagram of the overlap of significant CpGs (p < 10^−4^) in three species. Scatter plots DNA methylation aging in NMR, and mouse (C), and human (D). The highlighted CpGs are the loci that diverged in aging pattern between NMR and other species. E) DNA methylation aging in selected loci with divergent aging pattern in NMR, and other species.

**Supplementary Figure S10.**
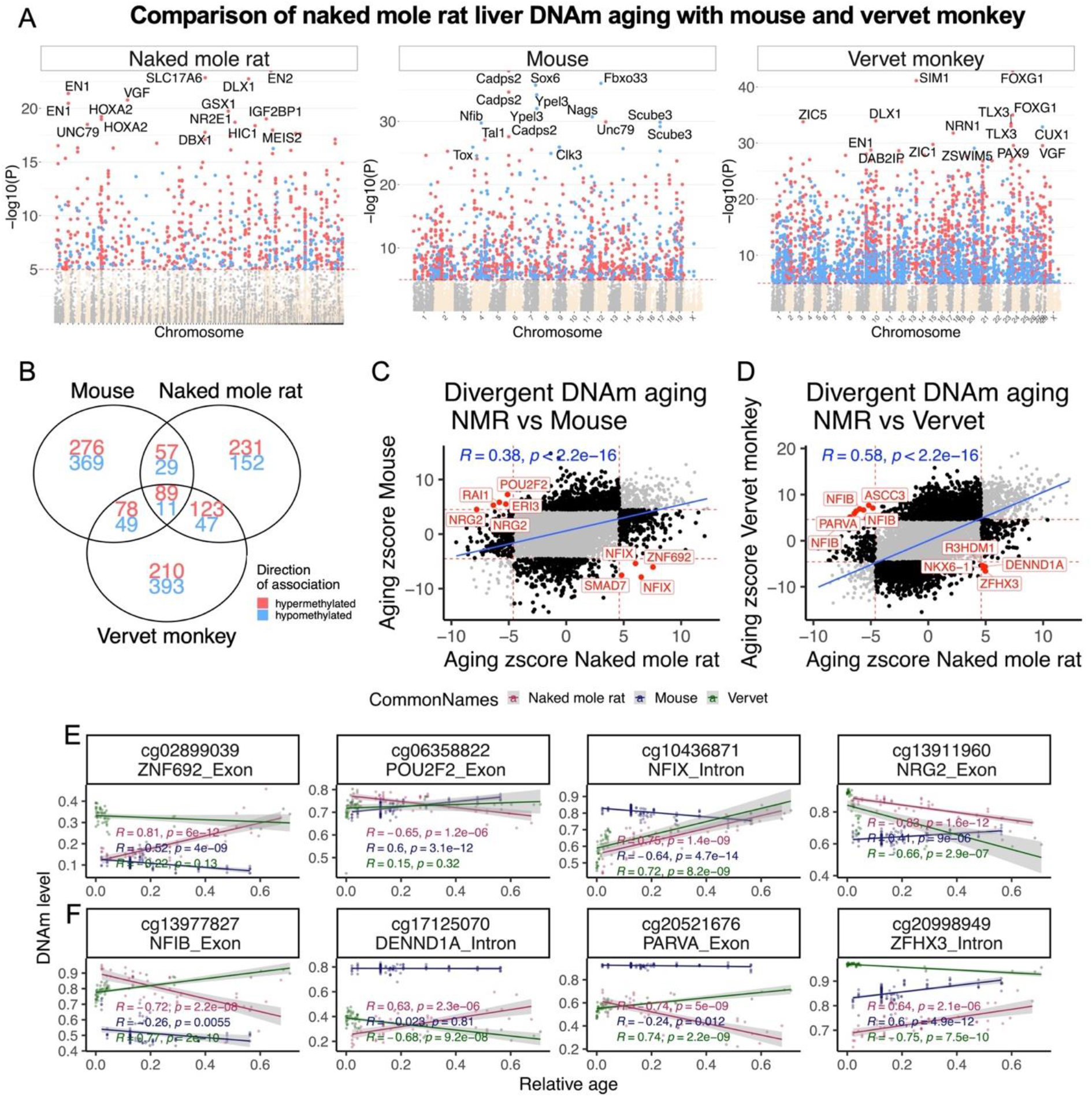
Naked mole-rat (NMR) liver has a distinct DNA methylation aging than mouse and vervet monkey. A) Manhattan plots of DNA methylation aging loci in NMR, vervet monkey and mouse. The red line in the Manhattan plot indicates p <10^−5^. The analysis was limited to 11,793 probes that aligned to orthologous genes in all three species. Sample sizes: NMR, 62; Mouse, 111; and vervet monkey, 48. The red line in the Manhattan plot indicates p <1e-5. B) Venn diagram of the overlap of the up to 1000 top CpGs (p < 10^−5^) in three species. Scatter plots DNA methylation aging in NMR, and mouse (C), and Human (D). The highlighted CpGs are the loci that diverged in aging pattern between NMR and other species. DNA methylation aging in selected loci with divergent aging pattern in NMR than mouse (E), or vervet monkey (F).

**Supplementary Figure S11.**
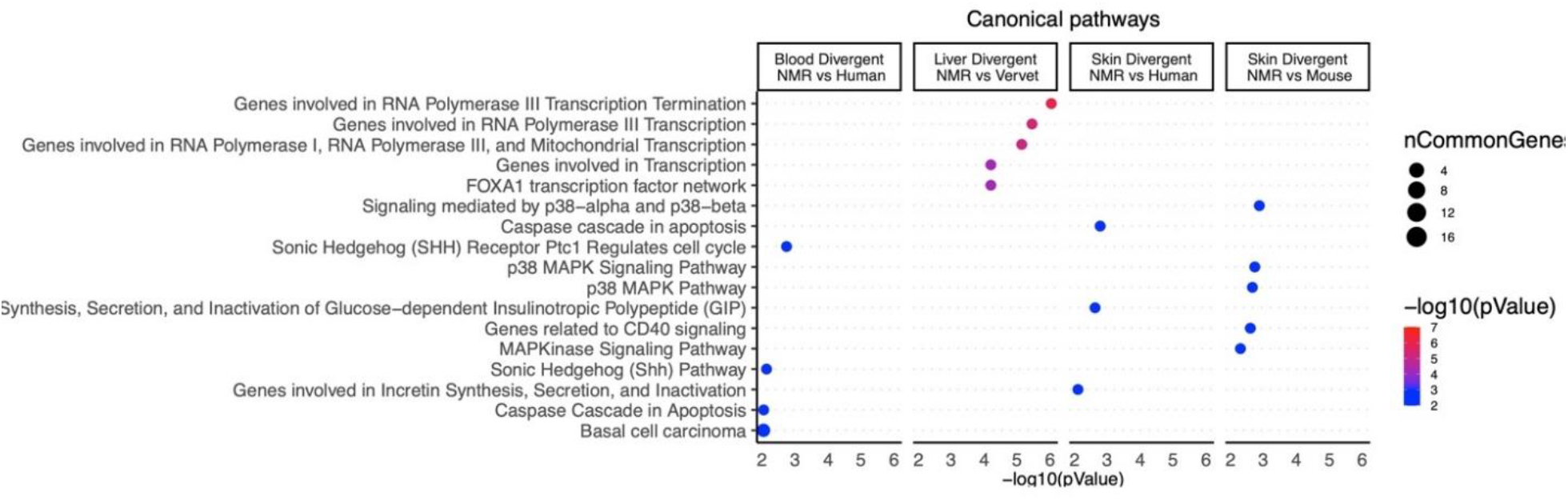
Enrichment results for CpGs with distinct aging patterns in the NMR. The columns correspond to different species comparisons between NMR, mouse, and primates (human or vervet monkey) in different tissues (blood, liver and skin). The analysis was done using the genomic region of enrichment annotation tool^60^. The gene level enrichment was done using GREAT analysis^60^ and human Hg19 background. The background probes were limited to 11,814 probes in skin and blood, or 11,793 probes in liver that were mapped to the same gene in three species.

**Supplementary Figure S12.**
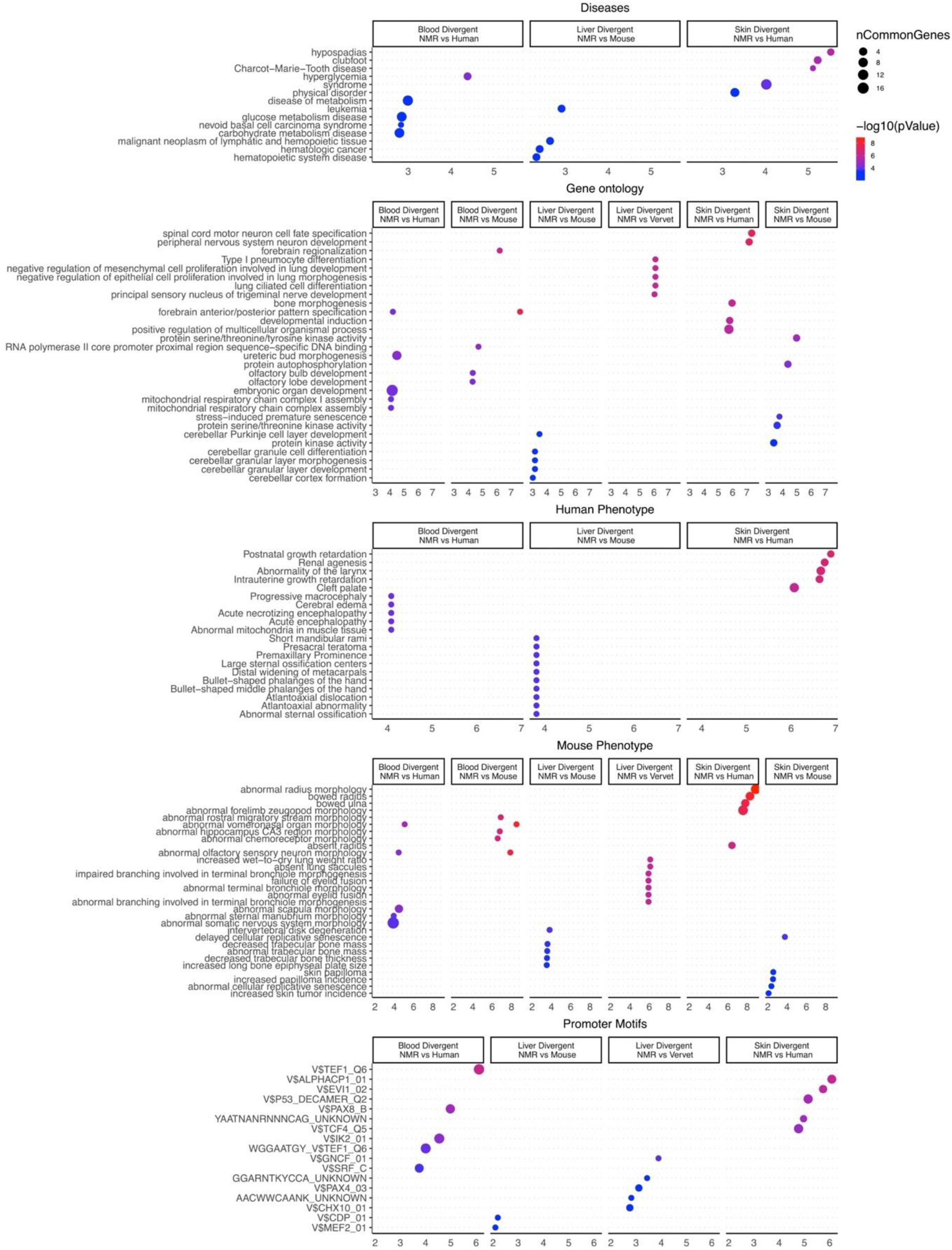
Gene set enrichment analysis of divergent DNA methylation patterns between NMR, human and mouse in liver and skin. The analysis was done using the genomic region of enrichment annotation tool^60^ The gene level enrichment was done using GREAT analysis^60^ and human Hg19 background. The background probes were limited to 11,814 probes that were mapped to the same gene in the NMR, human, and mouse genome. The top five enriched datasets from each category (Canonical pathways, diseases, gene ontology, human and mouse phenotypes, and promoter motifs) were selected and further filtered for significance at p < 0.01.

**Figure S13.**
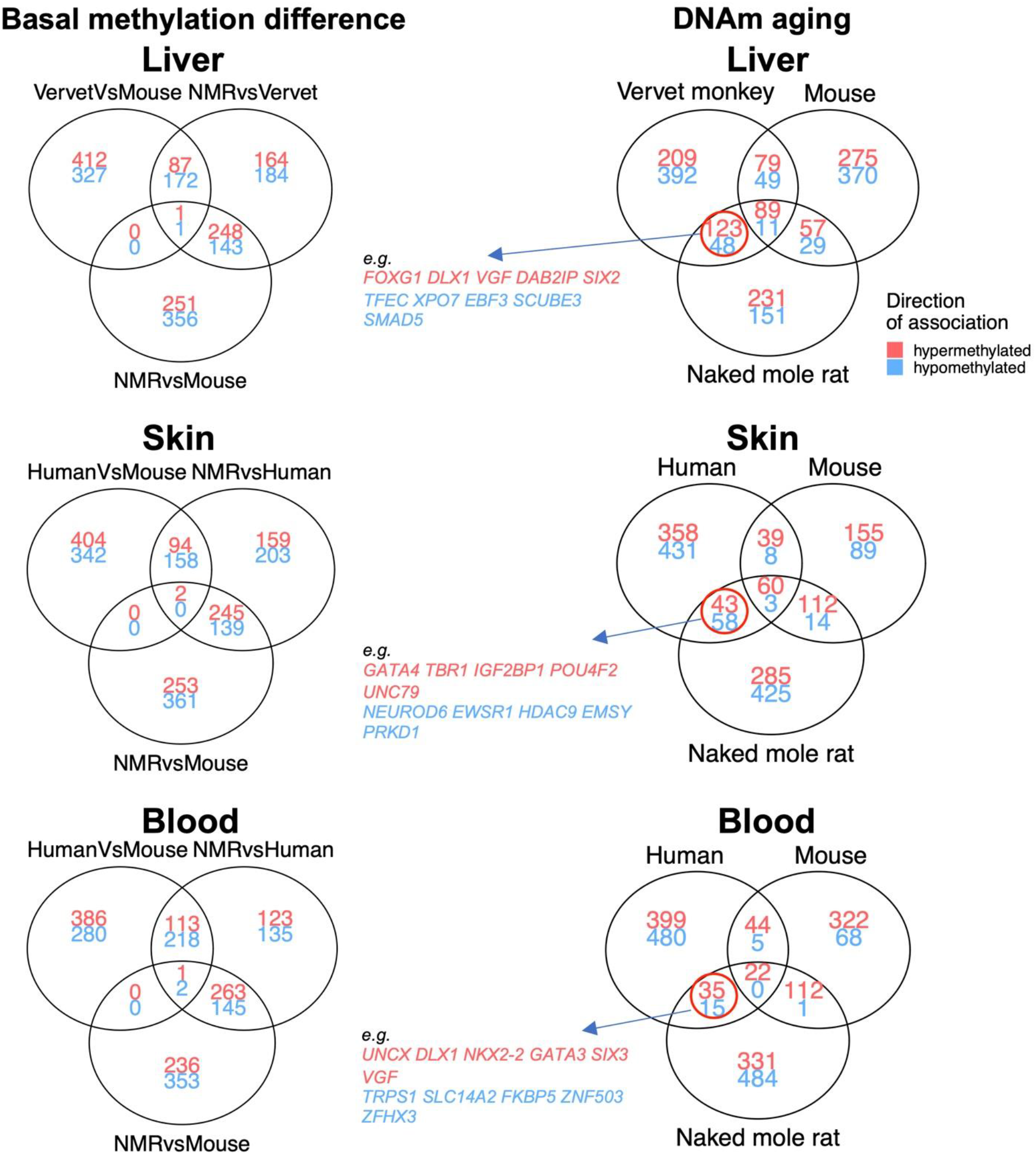
DNAm signatures that are shared between NMR and primates, but not with mouse. Venn diagrams represent the overlap of top CpGs in different EWAS analysis. The left column shows a pairwise basal (mean) DNAm difference between NMR, primates and mouse. The right column shows the overlap of top CpGs with DNAm aging between NMR, primates and mouse. We identified the changes that were shared between NMR and primates, but not in mouse. These CpGs are reported in Table S13.

**Figure S14.**
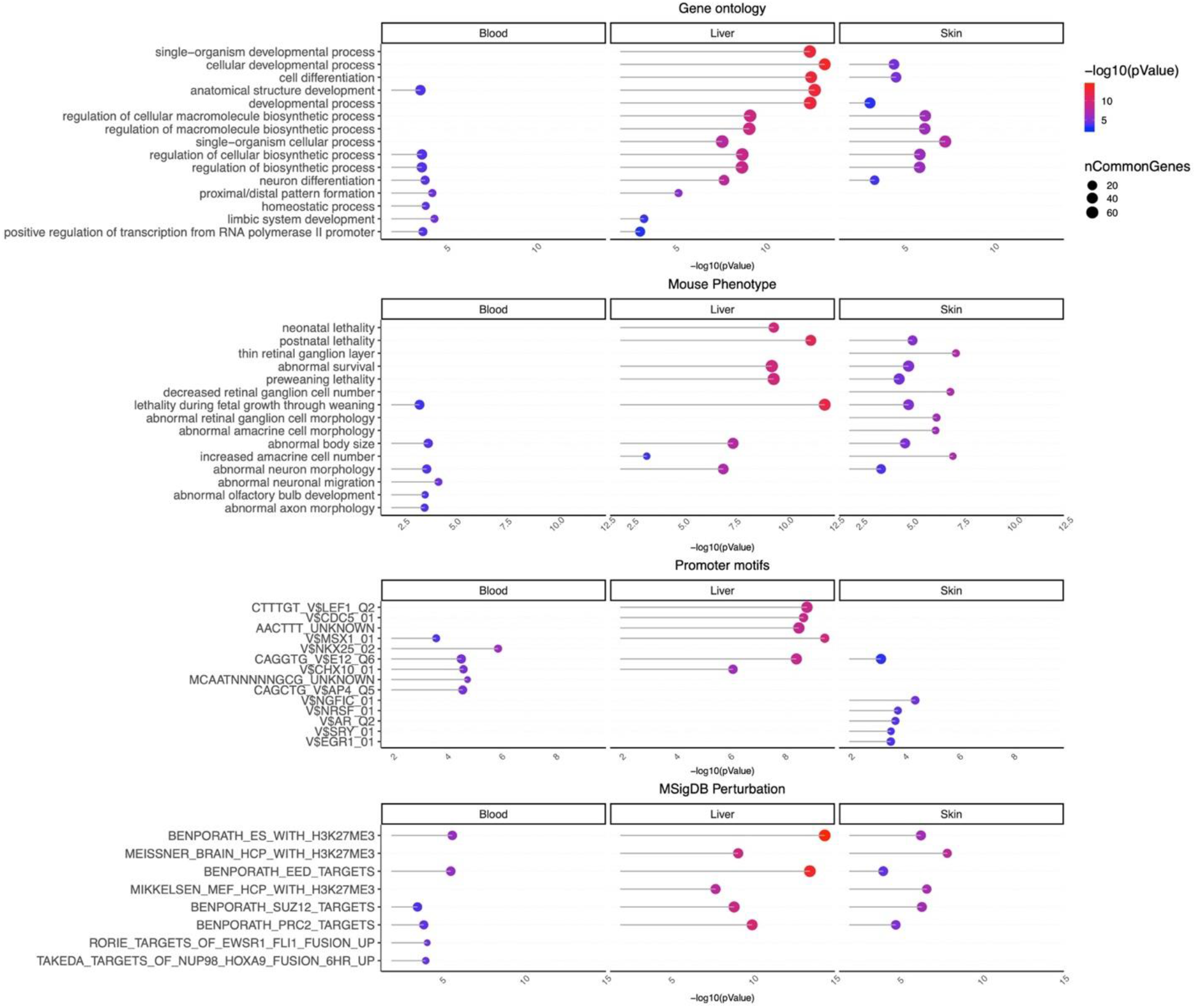
Enrichment analysis of the tissue specific DNAm signatures that are shared between NMR and primates, but not with mouse. The input includes the CpGs that changed with age only in NMR and primates in the same direction. The analysis was done using the genomic region of enrichment annotation tool^60^ The gene level enrichment was done using GREAT analysis^60^ and human Hg19 background. The background probes were limited to 11,814 probes that were mapped to the same gene in the NMR, human, and mouse genome. The top five enriched datasets from each category (Canonical pathways, diseases, gene ontology, human and mouse phenotypes, and promoter motifs) were selected and further filtered for significance at p < 1e-3.

